# Unraveling mitotic protein networks by 3D multiplexed epitope drug screening

**DOI:** 10.1101/231779

**Authors:** Lorenz Maier, Stefan Kallenberger, Katharina Jechow, Marcel Waschow, Roland Eils, Christian Conrad

**Affiliations:** German Cancer Research Center (DKFZ), Theoretical Bioinformatics Division, Im Neuenheimer Feld 267, 69120 Heidelberg, Germany; Heidelberg University, Institute for Pharmacy and Molecular Biology (IPMB), Bioinformatics and Functional Genomics Department, Im Neuenheimer Feld 364, 69120 Heidelberg, Germany

**Keywords:** Multiplexed immunostaining, 3D cell culture, protein-protein interactions, mitosis

## Abstract

Three-dimensional protein localization intricately determines the functional coordination of cellular processes. The complex spatial context of protein landscape has been assessed by multiplexed immunofluorescent staining^1–3^ or mass spectrometry^4^, applied to 2D cell culture with limited physiological relevance^5^ or tissue sections. Here, we present **3D SPECS**, an automated technology for **3D** Spatial characterization of **P**rotein **E**xpression **C**hanges by microscopic Screening. This workflow encompasses iterative antibody staining of proteins, high-content imaging, and machine learning based classification of mitotic states. This is followed by mapping of spatial protein localization into a spherical, cellular coordinate system, the basis used for model-based prediction of spatially resolved affinities of various mitotic proteins. As a proof-of-concept, we mapped twelve epitopes in 3D cultured epithelial breast spheroids and investigated the network effects of mitotic cancer drugs with known limited success in clinical trials^6–8^. Our approach reveals novel insights into spindle fragility and global chromatin stress, and predicts unknown interactions between proteins in specific mitotic pathways. **3D SPECS’s** ability to map potential drug targets by multiplexed immunofluorescence in 3D cell cultured models combined with our automized high content assay will inspire future functional protein expression and drug assays.

---

Iterative antibody labeling overcomes the spectral limit of total number of fluorescent antibodies that can be applied simultaneously to individual cells^1,3^. We successfully extend this technique of chemically bleached fluorescently labeled antibodies, to 3D cell cultured spheroids in Matrigel^9^ combined with drug treatment (see Supplementary Table 1). Our setup (**Fig. 1a**) uses confocal laser scanning microscopy together with automated pre-screen classification by machine learning and motorized in-built micro pipetting robot to identify and comprehensively stain mitotic phases. Naturally, mitotic cells self-organize, structure and typically segregate in different orientations, rendering a direct comparison of different data sets impossible. To overcome this limitation, we propose a novel representation named **SpheriCell** for spatial alignment of subcellular images. It infers a spherical coordinate system of the cellular space, using the spindle axis perpendicular to the metaphase plate to define the orientation of the vertical axis, while the metaphase plate defines the equatorial plane. Within this spherical coordinate system, protein concentrations are measured as 3D partitions in three symmetrical sets of spherical sectors and six shells. For enhanced visualization, mean values of 3D partitions are projected onto a two-dimensional longitudinal plane (**Fig. 1b**). Hence, we screened 6,272 image stacks, identified 1,217 mitotic events resulting in 284,778 mean intensity values of 3D partitions. We illustrate differences between tumorigenic and non-tumorigenic 3D cell lines in more physiological conditions still accessible by high-throughput screening^5^. The applied human epithelial MCF10 breast cancer progression model compares near-diploid cell line MCF10A^10^ forming polarized spheroids^11^ with the tumorigenic line MCF10CA, which bear activating mutations of *HRAS* and *PIK3CA*, amplified *MYC*, but wild-type *TP53*^10^.

**Figure 1.**
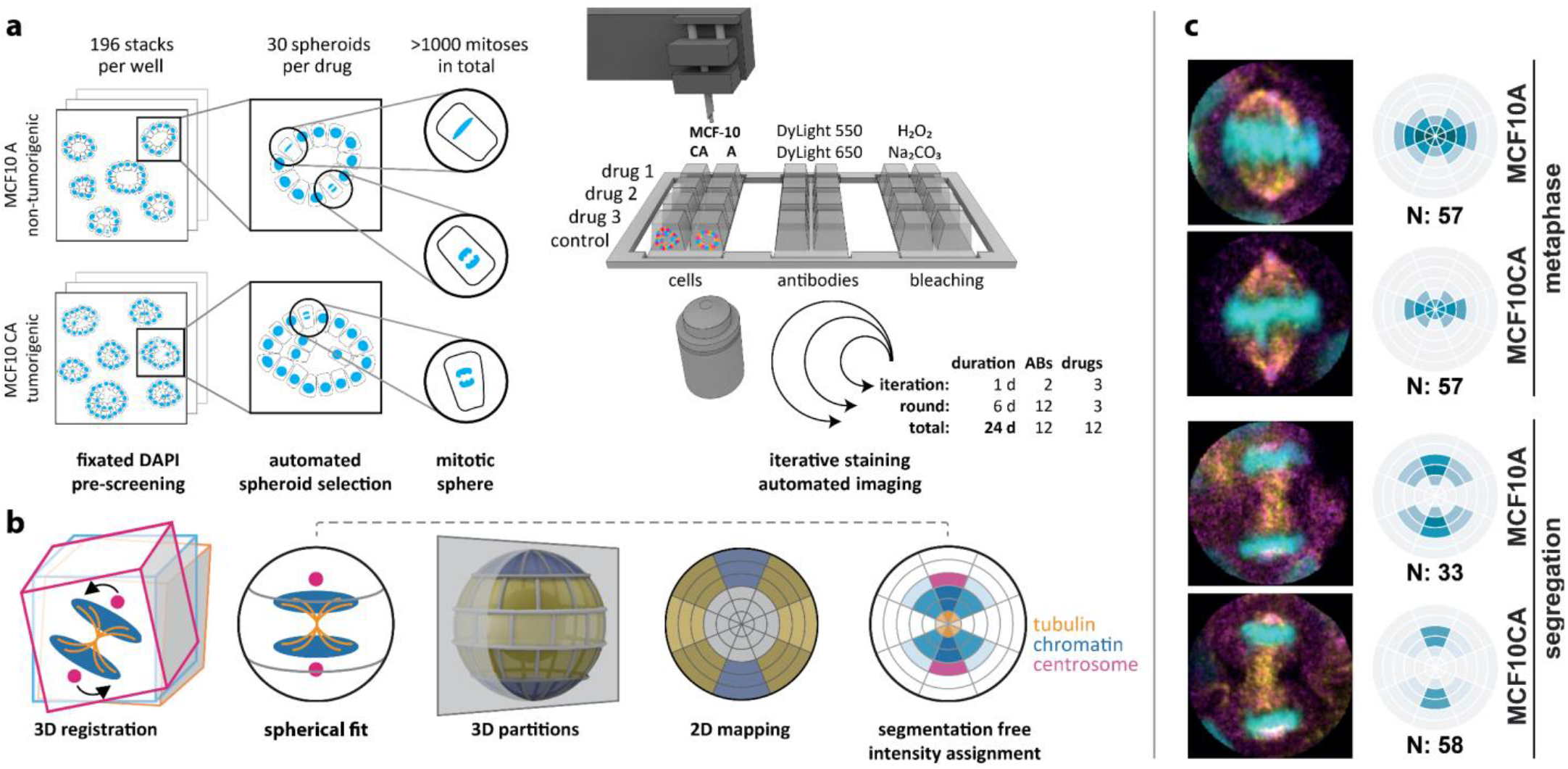
Workflow of iterative antibody labeling. (**a**) After 48h drug treatment, MCF10A and MCF10CA cells were fixed and DAPI stained. Pre-screening comprised 196 image stacks per well to automatically select 30 spheroids that each showed at least one mitosis. At each round, selected positions for three drug treatments plus control were stained, imaged, and bleached in six iterations with two antibodies each. Within 24 days, we acquired 3D stacks of twelve antibodies on twelve drug treatments and two cell lines. (b) **SpheriCell** visualization: Stacks were 3D registered and a sphere was fitted to each cell division area, which was partitioned into three symmetrical sets of spherical sectors and six shells. Spherical 3D localization can be visualized by a longitudinal cut resulting in a 2D polar grid that contains projected mean values of 3D partitions. Moreover, cell poles are not distinguishable, so the results are centrically symmetric. Localization of mitotic proteins can be intuitively determined from the 2D projected partitions as exemplified by tubulin, chromatin, and centrosomal regions. Color intensities reflect normalized, mean protein concentrations in each bin. (c) Example images and DAPI binning. Distinguished between MCF10A and MCF10CA, and metaphase and segregation spanning ana- and telophase. DAPI (cyan), γ-Tubulin (magenta), and β-Tubulin (yellow). Example images were rotated manually. ABs: antibodies; N: number of mitoses contributing to mean values.

Using **3D SPECS**, we first studied sub-cellular differences in protein localizations in the tumorigenic and non-tumorigenic 3D cell lines distinguishing between the cell cycle states metaphase and segregation. Here we assessed co-localization affinities of proteins and their preferred localization to subcellular compartments by our mathematical modeling approach. We thus unraveled network-wide treatment effects on mitotic spindle organization, spindle assembly checkpoint (SAC), and complementary cell fate indicators. The spindle assembly checkpoint (SAC) control includes e.g. the chromosome passenger complex (CPC) comprising BIRC5, Borealin, INCENP, and Aurora B^12^, which inhibit anaphase promoting complex (APC/C) most efficiently through mitotic checkpoint complex (MCC) containing BUB1 β (BUBR1)^13^. Failures in these checkpoints can lead to disruption of mitosis and subsequent autophagic or necrotic events.

We confirmed known localizations of cellular proteins and known mitotic checkpoints for untreated cells as described before^12–21^, supporting the utility of **3D SPECS** (**Fig. 2a**). MCF10CA staining patterns resembled those of MCF10A, showing a slightly reduced average DAPI signal due to increased cell size. Highest protein concentration increases were observed for γ-H2AX and Aurora *A*, contrasted by a reduction strongest for γ-Tubulin (**Fig. 2b**). Increased levels of γ-H2AX^16^, a marker for double strand breaks, may reflect higher chromosomal stress. Higher intensity levels of Aurora A, which is upregulated during mitosis and localizes mostly towards centrosomes^14^, in MCF10CA is consistent with previously described effects upon activation of Raf-1^22^, downstream of the oncogenic RAS pathway.

**Figure 2.**
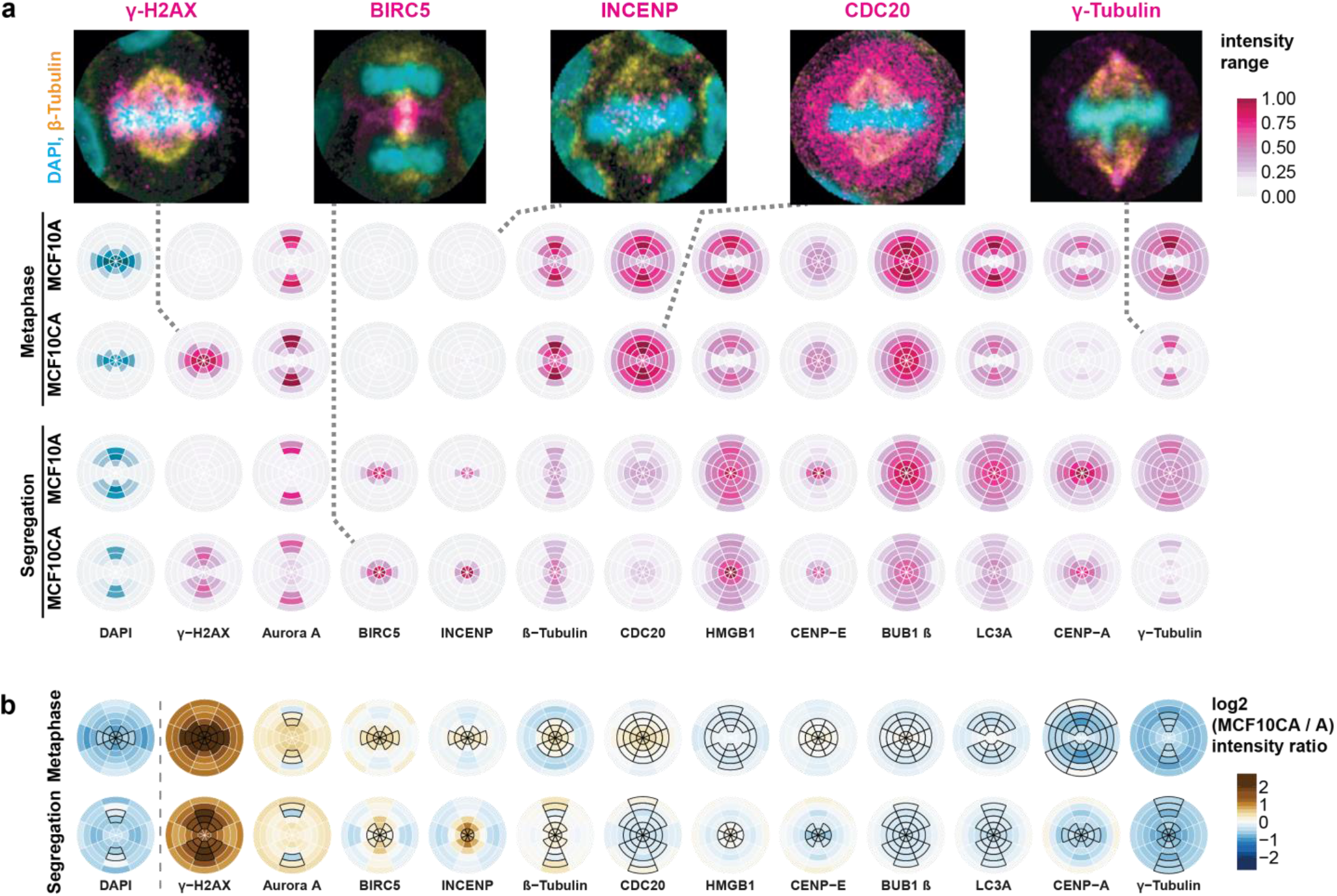
Localization and intensity changes of twelve antibody stainings of untreated MCF10A and MCF10CA cells during metaphase and segregation comprising ana- and telophase. SpheriCell plots depict mean intensity values across all imaging rounds, ordered by decreasing difference between MCF10CA and MCF10A cells. (**a**) Localization of epitopes. Intensity ranges were specific to the antibody and are shown normalized between 0 and 1, effectively across all values of a column in the figure. Distribution patterns generally reflect the localization of individual proteins described before^12–21^. Dashed lines connect SpheriCell plots with example images of antibody stainings (magenta), DAPI (cyan), and β-Tubulin (yellow). (**b**) MCF10CA shows altered intensity patterns compared to MCF10A. SpheriCell plots depict differences of log2 transformed fluorescence intensity of MCF10CA and MCF10A [log2(CA) – log2(A)] for metaphase and segregation, in decreasing order. Black framed partitions indicate intensity distributions in untreated control images. LC3A: microtubule-associated proteins 1A/1B light chain 3A.

We then analyzed the effects of twelve targeted inhibitors on mitotic proteins of dividing MCF10A and MCF10CA cells (**Fig. 3a,b**), specifically on protein concentrations and preferred localizations. To compare spatial distribution patterns of protein intensities, in each cell, 18 subcellular compartments were defined by a combination of six eccentricity shells with three orientations relative to the division plane. In analogy to calculating a center of mass, we specified measures of spatial intensity distributions (**Supplementary Note 1**). **Fig. 3a** visualizes significant concentration fold changes and spatial changes in eccentricity and orientation compared between cell lines, mitotic phases and for inhibitors relative to controls. Notably, comparisons between cell lines showed more pronounced effects on concentration fold changes than on spatial distributions. MCF10CA cells showed significantly higher γ-H2AX and Aurora A concentrations, but lower γ-Tubulin concentrations. Changes in eccentricity and orientation of the localization pattern could be observed in MCF10CA relative to MCF10A cells mostly during metaphase. Obviously, as indicated for the comparison between segregation and metaphase, the mitotic phase strongly influenced the spatial distributions of most observed proteins. Changes in the distribution pattern for DAPI and γ-H2AX reflect the movement of the nucleus towards the cell division axis and to higher eccentricity while the other proteins move closer to the cell division plane.

**Figure 3.**
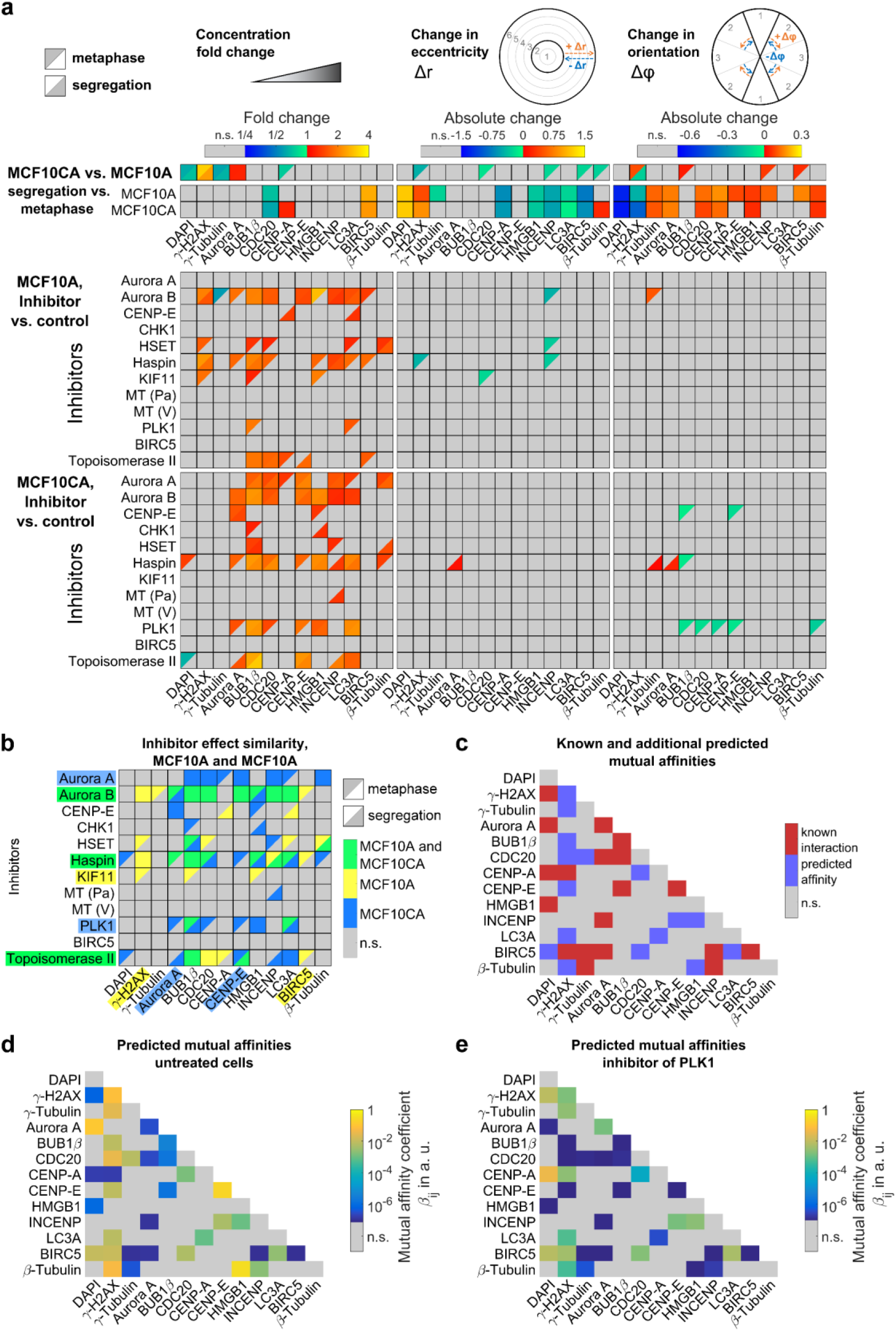
Inhibitor effects and estimates of affinities between proteins. (**a**) Concentration fold changes and localization changes, quantified as changes in eccentricity and orientation of localization patterns for comparisons between cell lines (MCF10CA vs. MCF10A), mitotic phases (segregation vs. metaphase) and inhibitor treatments (inhibitor vs. control). The left column shows color-coded fold changes in average concentrations (total intensities normalized by cell volumes) for DAPI and antibody stainings on a logarithmic scale. In the center column, eccentricity changes for intensity distributions in subcellular compartments were visualized. Positive values describe a movement to the periphery while negative values represent a movement to the center of the cell. Similarly, in the right column, changes in angular orientation of intensity distributions were visualized. Positive values describe a movement towards the plane perpendicular to the cell division axis while negative values describe a movement towards the cell division axis. In cases of significant differences to negative controls (t-test with p<0.05, followed by Bonferroni multiple testing correction for 54 comparisons in each column), fold changes relative to negative controls are indicated by colors (n. s., not significant). Values indicated separately for metaphase (upper left triangles) and segregation (lower right triangles) in (a) and (b). (**b**) Overlay of significant effects in MCF10A and MCF10CA cells, MCF10A only, or MCF10CA cells only. Color highlighted proteins denote predominant effects per row or column. (c) Known protein-protein interactions from Ingenuity Pathway Analysis overlaid with additional predicted mutual affinities between measured proteins (see **Supplementary Note 1**). For affinities to subcellular compartments, see **Supplementary Fig. 1** (**d**) Estimates of mutual affinities between measured proteins for untreated cells. (**e**) Estimated mutual affinities between measured proteins after treatment with an inhibitor of PLK1.

During segregation, relative to metaphase, both cell lines showed a known decrease in CDC20 concentration^23^ and elevated BIRC5 concentrations, whereas the CENP-A was only increased in MCF10A cells. Inhibitors mostly increased the concentrations of several proteins but hardly affected their spatial distributions (**Fig. 3b**). Here, the inhibitions of master regulator AURKB^12^ and also Haspin known to be implicated in Aurora B positioning^12^ showed a prominent effect across nearly all proteins in MCF10A as well as MCF10CA cells. Moreover, we detected broad effects of increased DNA damage by Topoisomerase II poisoning^24^. Interestingly, MCF10CA cells appeared to be more sensitive to mitotic spindle interference, reflected by effects on Aurora A and CENP-E. Concordantly, inhibition of Aurora A affected more proteins in MCF10CA spheroids compared to MCF10A. MCF10CA showed more pronounced effects of PLK1 inhibition. Contrarily, although KIF11 (Eg5) and KIFC1 (HSET) facilitate separation and clustering of centrosomes^25^, the effects due to inhibition of KIF11 were restricted to MCF10A cells. High natural levels of γ-H2AX intensity in MCF10CA were not increased by our treatments as observed in MCF10A. Also, BIRC5 concentration was affected in MCF10A but not in MCF10CA cells. In contrast to effects on concentrations, the eccentricity and the orientation of distribution patterns were less affected by inhibitions except for Haspin and PLK1 being two notable exceptions.

To study intracellular distributions and localized interactions between proteins, we developed a non-linear model that was calibrated with our partitioned mitotic protein intensity data (**Supplementary Note 1**). To capture interactions between proteins in a simplified manner, the model describes concentrations at steady state for monomers, homo- and heterodimers of all proteins bound to subcellular compartments with first and second order interactions. Affinities of every protein to subcellular compartments, and affinities between proteins were estimated by model fitting. To predict new affinities between proteins, we started by fitting a model of interactions from literature in *Ingenuity* Pathway Analysis (IPA) regarded as ground truth. This initial model was fitted to our untreated control cells. Then, by sequential forward selection, new mutual affinities between proteins were additionally included in the model and pertained if model fits were significantly improved, based on likelihood-ratio testing. **Fig. 3c** visualizes the 19 known interactions overlaid with all 16 additionally predicted mutual affinities. Interestingly, colocalization affinities between species do not necessarily imply functional relations and pathway interactions do not require high affinities. For example, we identified known interactions of γ-Tubulin with CDC20^26^ as well as known DNA-binding of BIRC5^12^ (**Fig. 3c**). While we triggered DNA damage pathways with Topoisomerase II poisoning and inhibition of CHK1^27^, indications of cell fate could be inferred from double strand break marker γ-H2AX^16^, necrosis associated HMGB1^17^, and autophagic vesicle marker MAP1LC3A (LC3)^18^. BIRC5 was predicted by the model to interact with LC3, which links mitotic surveillance and autophagy pathways^18^. The model predicted interactions of γ-H2AX, γ-Tubulin and β-Tubulin with several other proteins, which might indicate indirect interactions with the mitotic spindle or the cytoskeleton. Coefficients describing mutual affinities between proteins are depicted in **Fig. 3d**. Importantly, the fraction of a protein that is localized to a mitotic bin due to mutual interactions with other proteins does not only depend on affinity coefficients but may require high affinities of interacting species to the respective mitotic bin. Estimated fractions of proteins recruited to subcellular compartments due to mutual interactions with other proteins, as well as estimated affinities to subcellular compartments are shown in **Supplementary Fig. 1**. The highest values of mutual affinities with other proteins were found between CENP-E molecules as described earlier^28^ and for the newly predicted binding of β-Tubulin to HMGB1 (**Fig. 3d**).

We next inspected changes in mutual affinities between mitotic proteins and their affinities to subcellular compartments upon drug treatment. To this end, the model with known and additionally predicted interactions was fitted to datasets from cells treated with inhibitors to estimate mutual affinities between proteins and to subcellular compartments. In **Fig. 3e**, estimated affinity coefficients for PLK1 as an exemplary inhibitor are visualized. Inhibition of PLK1 affected the localization patterns of several proteins and caused a strong specific shift in mutual affinities among several studied proteins in comparison to untreated cells (**Fig. 3d,e**). In those cases, many estimates of mutual affinities decreased, indicated by changes to blue color code. Contrarily, affinities to subcellular compartments during metaphase and segregation showed almost no differences to untreated cells (**Supplementary Fig. 1e-h**). It is tempting to speculate that effects of PLK1 inhibition are mediated through its involvement in spindle network formation^29^. Specifically, predicted affinity of β-Tubulin to γ-H2AX, HMGB1 and INCENP decreased, and chromosome affinity of BIRC5 appears to be reduced by inhibition of PLK1, whereas the predicted affinity of INCENP to HMGB1 is increased (**Fig. 3e**). Reduced chromosome affinity of BIRC5 after PLK1 inhibition is in accordance with the finding that phosphorylation of BIRC5 by PLK1 is required for a proper chromosome alignment during mitosis^12^.

To summarize, we present **3D SPECS**, a high-content screening assay employing automated iterative antibody labeling in 3D cell cultures. It allowed us to compare system-wide interactions between twelve proteins of two cell lines in two mitotic phases, upon twelve individual treatments. High automation comprises detection of mitoses, iterative staining and imaging, 3D partitioning, modeling and visualization using **SpheriCell**, a novel approach that does not require antibody image segmentation, nor alignment of cell division in 3D. This explorative approach recapitulated *prior* knowledge on proteins involved in mitosis and allowed the generation of novel hypotheses in mitotic pathway signaling. Most prominently, we discovered up-regulation of γ-H2AX in tumorigenic MCF10CA cells compared to MCF10A. Further, γ-H2AX has a higher sensitivity to interference in MCF10A, which in turn appears to have a more robust spindle apparatus. Our novel combined imaging and mathematical modeling approach allowed us to disentangle inhibitor-mediated protein localization and binding affinity changes and showed that changes in affinities between proteins (**Fig. 3d,e**) were more pronounced than changes in individual protein localizations (**Supplementary Fig. 2d,e**). As an example, we focused on the measured inhibitions of PLK1 activity, which is responsible for establishing the mitotic spindle and which is frequently hyper-activated in cancer^30^. Subsequent reduction in chromatin affinity of BIRC5 could be explained by its dependency on PLK1 phosphorylation^12^, most likely intertwined with CPC function.

Our method can be readily extended to directly determine the activity of proteins by phospho-specific antibodies. For a more fine-grained assessment of protein localization additional nuclear or membrane staining can be easily integrated into **3D SPECS**. The **SpheriCell** approach that renders intuitively simple and comprehensive visualization of protein localization in cell division can also be amended by including polar landmarks of non-dividing cells. Taken together we have demonstrated **3D SPECS** as a novel workflow unraveling thus unprecedented levels of details in changes of protein localization and interaction upon drug treatment of three-dimensional cell cultures.

## Acknowledgements

We thank Sabine Aschenbrenner for support with lab techniques, Siegfried Winkler, Leo Burger, and Helmuth Schaar for microscopy hardware, Antonio Politi for imaging advice and ZEN black macro interface, Maria Maier for assistance with 3D renderings, Christian Dietz for continued development of KNIME image processing, and Clarissa Liesche and Joël Beaudoin for critical comments. The authors acknowledge support by the state of Baden-Württemberg through bwHPC.

## Author contributions

L.M. and C.C conceived the experiments and subcellular visualization strategy. L.M., K.J. and M.W. established antibody staining and drug treatment protocols. L.M. developed automated iterative staining workflow, conducted experiments, and image analysis. S.K. developed the interaction model and performed statistical tests. C.C. and R.E. supervised this project. L.M., S.K., C.C, and R.E. wrote the manuscript. All authors commented on the manuscript.

## Online Methods

Mitotic proteins were assessed after 48h drug treatment by iterative immunofluorescence labeling. The antibodies were either labeled with one of DyLight 550 / Cy3 or DyLight 650 / Cy5. Both types of dyes could be used interchangeably in terms of excitation and emission spectra.

While Matrigel is essential for acinar growth of spheroids^11^, it also dissolves quickly when the bleaching solution is applied. Therefore, we have used DyLight instead of Cy or Alexa^31^ labeled antibodies, as they bleach much faster and have a very strong fluorescence signal nevertheless. Applying the bleaching solution significantly longer than 5 minutes at a time typically dissolved the Matrigel carrying the spheroids.

All treatments were imaged at 30 spheroids that showed at least one mitosis each. 196 stacks per well, eight wells per round and four rounds resulted in 6272 image stacks with 21 slices each that were automatically pre-screened for mitotic events. Iterative high resolution images of 30 positions per well in eight wells in each of four rounds totaled in 960 identified spheroids that were imaged each with 31 slices, after staining and after bleaching in six iterations.

For analysis and visualization, every mitosis was aligned along its division plane for a spherical neighborhood that contains the cell division in equatorial axis (see Fig. 1b).

### 3D cell culture and drug treatment

Human mammary epithelial MCF10A pBabePuro cells were kindly obtained from Zev Gartner Lab; MCF10CA1d.cl1 (MCF10CA) cells from Karmanos Cancer Institute. Eight well Lab-Tek Chambered Coverglass slides (Sigma 155411) were treated with 2 M NaOH for 20 min and rinsed twice for 10 min with MilliQ water. Ten μl Matrigel (growth factor reduced, phenol red-free, Corning 356231) per well was added on ice with pipette tips pre-cooled to -20 °C. MCF10A and CA cells were seeded with 2% Matrigel in Growth Medium overnight. Growth medium was adapted from Debnath et al.^32^ and is based on DMEM/F12 (no phenol red, Gibco 21041-33), with 5% Horse Serum (Gibco 16050-122), 20 ng/ml EGF (Sigma E9644-.2MG), 0.5 mg/ml Hydrocortisone (Sigma H0888-1g), 100 ng/ml Cholera Toxin (Sigma C8052-1MG), 10 μg/ml Insulin (Life Technologies 12585014). For the inhibition experiments, the cells were treated for 48h at one day after seeding.

### Inhibitors

Drugs, suppliers, and concentrations used were Barasertib (Aurora B inhibitor; alternative name AZD1152-HQPS; SelleckChem S1147; 1.11 nM); CHR-6494 (Haspin inhibitor; MedChem Express HY-15217; 500 nM); CW069 (HSET inhibitor; SelleckChem S7336; 25.0 μM); Etoposide (Topoisomerase II inhibitor; SelleckChem S1225; 333 nM); GSK461364 (PLK1 inhibitor; SelleckChem S2193; 2.20 nM); GSK923295 (CENP-E inhibitor; SelleckChem S7090; 3.20 nM); Ispinesib (KIF11 inhibitor; alternative name SB-715992; SelleckChem S1452; 1.70 nM); MK-5108 (Aurora A inhibitor; alternative name VX-689; SelleckChem S2770; 0.576 nM); MK-8776 (CHK1 inhibitor; alternative name SCH 900776; SelleckChem S2735; 9.00 nM); Paclitaxel (microtubule inhibitor; SelleckChem S1150; 2.67 nM); Vinblastine (microtubule inhibitor; Sigma V1377; 2.40 nM); and YM155 (BIRC5 inhibitor; SelleckChem S1130; 0.540 nM).

### Antibodies and labeling kits

Antibodies were conjugated with DyLight 550 and 650 Microscale labeling kits per supplier reference manual (Sigma, 84531 and 84536, respectively) unless otherwise stated. Antibody targets, dilutions, supplier, and conjugation method in iterative staining order were CENP-E (1:400; Abnova MAB1924; conjugated DyLight 550); BubR1 (1:600; Thermo Fisher MA5-16036; pre-conjugated with DyLight 650); beta-Tubulin (1:5000; Abcam ab11309; pre-conjugated with Cy3); CDC20 (1:400; Bethyl A301-179A; conjugated DyLight 550); gamma-Tubulin (1:12000; Abcam ab176404; pre-conjugated with Cy3); LC3A, microtubule-associated proteins 1A/1B light chain 3A (1:400; Novus NB100-2331; conjugated DyLight 650); BIRC5 (1:1000; Abcam ab176402; pre-conjugated with Cy3) INCENP (1:1000; Thermo Fisher MA5-17100; conjugated DyLight 650); Aurora A (1:6000; Abcam ab176375; pre-conjugated with Cy3); CENP-A (1:500; Abnova PAB18324; conjugated DyLight 650); HMGB1 (1:3000; Abcam ab176398; pre-conjugated with Cy3); H2AX (1:2500; Cell Signaling 9718BF; conjugated DyLight 650).

### Iterative antibody labeling

Cell fixation was based on a protocol from Debnath et al.^32^, with 1.85% formaldehyde solution (Sigma 252549) added to the medium for 10 minutes. Cells were rinsed twice with PBS and permeabilized for 10 min at RT with 0.5% TX-100 pre-chilled to 8 °C, washed three times with PBS-glycine (130 mM NaCl, 7 mM Na_2_HPO_4_, 3.5 mM NaH_2_PO_4_, 100 mM glycine) for 10 minutes, and blocked overnight at RT in a blocking solution consisting of IF-wash solution^33^ (130 mM NaCl, 7 mM Na_2_HPO_4_, 3.5 mM NaH_2_PO_4_, 7.7 mM NaN_3_, 0.1% BSA, 0.2% Triton X-100, 0.05% Tween 20) with 10% goat serum (Sigma G9023-10ML) and 1:1000 DAPI (Sigma D8417-1MG), inside an opaque EMBL microscope incubation chamber. For each iteration, two antibodies were diluted in freshly prepared blocking solution and stored in a slide within a 4x LabTek holder (EMBLEM LTT-01 and LTT-02). They were automatically pipetted into the wells by a peristaltic pump of the ProCellcare 5030 system (ProDesign) and incubated for 3h, washed twice with IF-wash for 5 min and three times with PBS for 5 min. After imaging, freshly prepared H_2_O_2_ bleaching solution^34^ containing 3% H_2_O_2_ (AppliChem, Cat. No. 121076) and 0.1M Na_2_CO_3_/NaHCO_3_ buffer at pH ≈ 10 was stored in another LabTek. It was automatically applied for 5 minutes and washed twice with PBS for 5 minutes. Standard incubator light source was switched on during bleaching with Energenie EG-PM2. Pipetting positions were planned with Zeiss Zen blue (www.zeiss.com/zen) and pipetting workflow was implemented in LabView (www.ni.com/labview).

### Pre-screen

During blocking, slides were imaged with a Yokogawa CSU-X1 spinning disc unit attached to a Zeiss Observer Z1 inside an EMBL incubation chamber. 196 image stacks of 401.6 × 400 × 60 μm were taken per well with a plan-apochromat 20x/0.8 NA objective. Stack slices had 1004×1002 pixels, step size was 3 μm and exposure 40 ms. Candidate mitotic positions were detected via their DAPI signal by a custom KNIME workflow and selected or expanded manually if necessary. The automatic selection excludes monolayer slices and uses a supervised tree ensemble classifier (comparable to a random forest). For each treatment and cell line, 30 positions of spheroids with at least one mitosis each were selected for imaging during the iterative staining workflow.

### Acquisition of iterative staining images

After each round of bleaching or staining, spheroids were automatically imaged with a laser scanning confocal Zeiss LSM 780 connected to the same Axio Observer as the spinning disc unit. Stack dimensions were 106.07 × 106.07 × 60 μm, with 512 × 512 pixels per slice and 2 μm Z steps. Objective was plan-apochromat 20x/0.8 NA, pixel dwell time 3.15 μs, and pinhole 32 μm. Emission spectra were taken at 410 – 489 nm (DAPI), 560 – 586, 586 – 612, and 612 – 630 (three parts of Cy3 / DyLight 550), and 638 – 758 nm (Cy5 / DyLight 650).

### Image processing

Splitting the emission spectrum from 560 to 630 nm in three parts allowed for post-acquisition exposure correction. Only for BIRC5 it was necessary to exclude the strongest emission channel from the labeling image. All remaining split channels were averaged. Mitotic DAPI signal was segmented in 3D by a customized region growing algorithm^35^ with manual seeds and borders to closely neighboring nuclei, especially in Z-direction. Segmented areas were manually annotated with their mitotic phase and ana- / telophases joined. Assignment of segregating split chromatin regions to a single dividing cell was verified with β-Tubulin staining. Registration of consecutive stacks per imaging position used subpixel alignment of Fiji^36^ plugin Correct 3D Drift^37^ followed by MultiStackReg^38^ with scaled rotation. Registration of the stacks was verified manually. For the spherical neighborhood, images were interpolated linearly in Z to match the X/Y pixel dimensions. Spherical segment angles were generated by Recursive Zonal Equal Area Sphere Partitioning Toolbox^39^ in MATLAB (www.mathworks.com) for 180 areas. A custom R script was created to join the areas to segments and to fit them in size and orientation to the individual mitoses, which were identified with 3D ellipsoid fit of the 3D ImageJ suite^40^. Metaphases could use those values as-is, but the size of segregating cells was overestimated by the ellipsoid fit and replaced by the centroid distances of their individual chromatin regions. Their 3D orientation used the average of the first two eigenvectors and the normalized centroid to centroid vector as third. The six spherical neighborhood shells grow linearly in their radius from the mitotic center, and the inner four span the identified nucleus area. Missing cells due to loss of Matrigel or failed registration were removed from further analysis. All image processing steps were embedded in a KNIME^35^ workflow.

### Visualization

Antibody intensities are depicted as color-coded mean values for three distinct segment classes (phi levels, **Fig. 1b**). Decreased intensity dependent on imaging depth was corrected by the mean DAPI intensity within the individual spherical neighborhood and mean DAPI intensity per well in pre-screen images after z-projection. To avoid an artificial increase in background signal of antibodies, DAPI intensities below a minimum threshold were excluded. Highlighting of partitions was determined by the control intensity over all rounds. Data was visualized with the packages ggplot2^41^, EBImage^42^, and shiny (shiny.rstudio.com) for R (www.r-project.org).

### Statistics and mathematical modeling

To test for significance of comparisons between controls and inhibitor treatments, we applied two-sample t-tests. For comparisons between two groups, intensity values for all subcellular compartments were collected within each group. In total, 54 comparisons were made between measurements of each protein. Therefore, t-tests with p<0.05/54 were regarded as significant (Bonferroni correction for multiple testing). The interaction model was implemented in MATLAB. For model calibration, we applied the solver lsqnonlin using the trust-region-reflective algorithm. Details on the model formulation and fitting can be found in **Supplementary Note 1**.

### Interpretation

Known interactions form literature were generated through the use of QIAGEN’s Ingenuity^®^ Pathway Analysis (IPA^®^, QIAGEN Redwood City, www.qiagen.com/ingenuity). They are supported by at least one reference from the literature, from a textbook, or from canonical information stored in the Ingenuity Pathways Knowledge Base.

### Data and code availability

Spherical neighborhoods, their image sources, and custom software code is available upon request.

## Supplementary Material

**Supplementary Figure 1.**
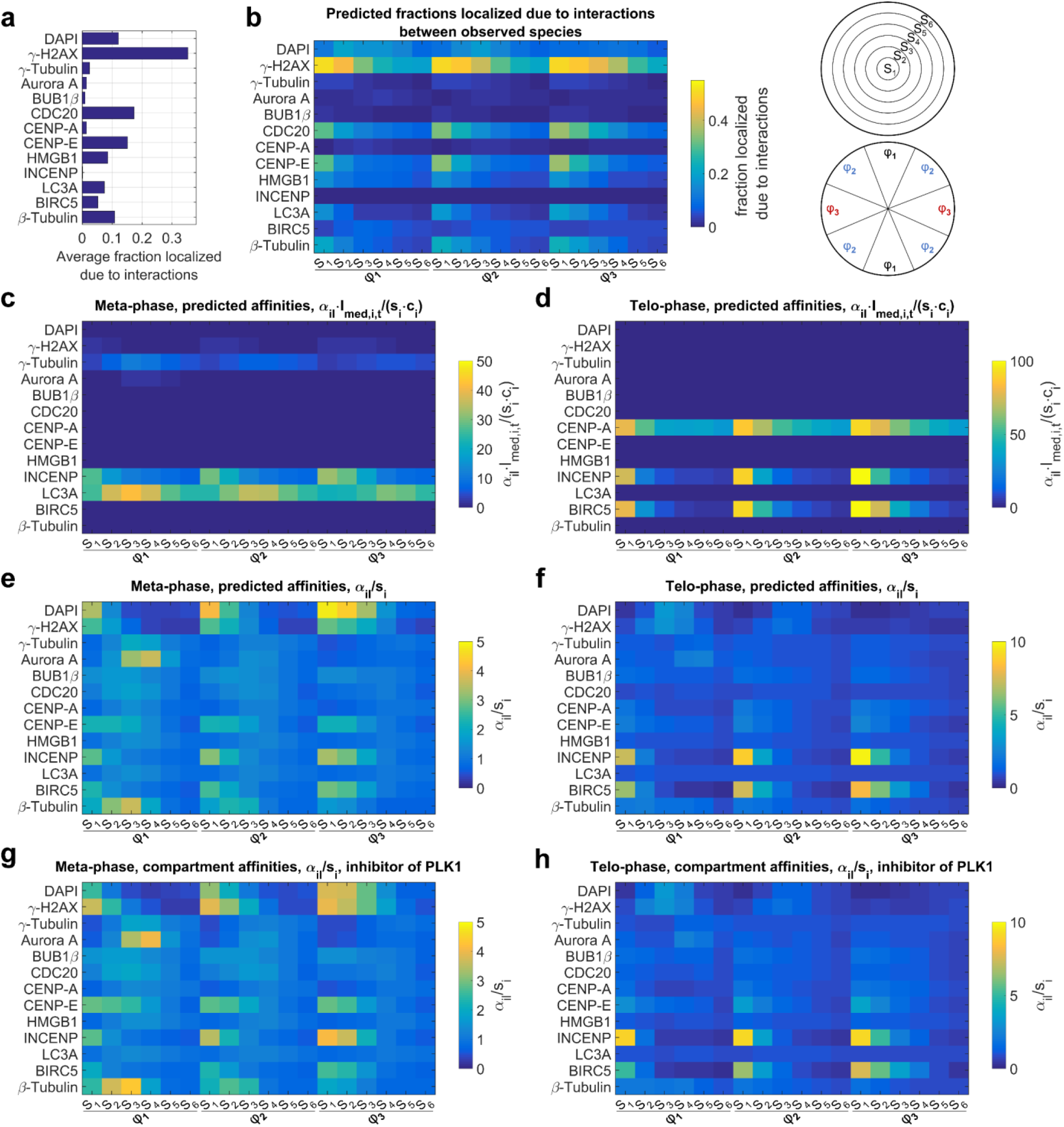
Predicted localization affinities. (**a**) Predicted fractions of proteins recruited to subcellular compartments due to mutual affinities between proteins. (**b**) Predicted fractions of proteins recruited due to mutual affinities between proteins, distinguished by eccentricity intervals (S_1_ to S_6_) and orientations (φ_1_ to φ_6_) as indicated by schematic maps. (**c**) Affinity estimates 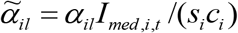, before dividing by scaling factor estimates, during metaphase or segregation obtained by model fitting to dataset from untreated cells (see **Supplementary Note 1** for details). (**d**) Rescaled untreated localization affinities *α_il_* /*s_i_* during metaphase or segregation. (**e**) Rescaled localization affinities *α_il_* /*s* upon treatment with PLK1.

**Supplementary Figure 2.**
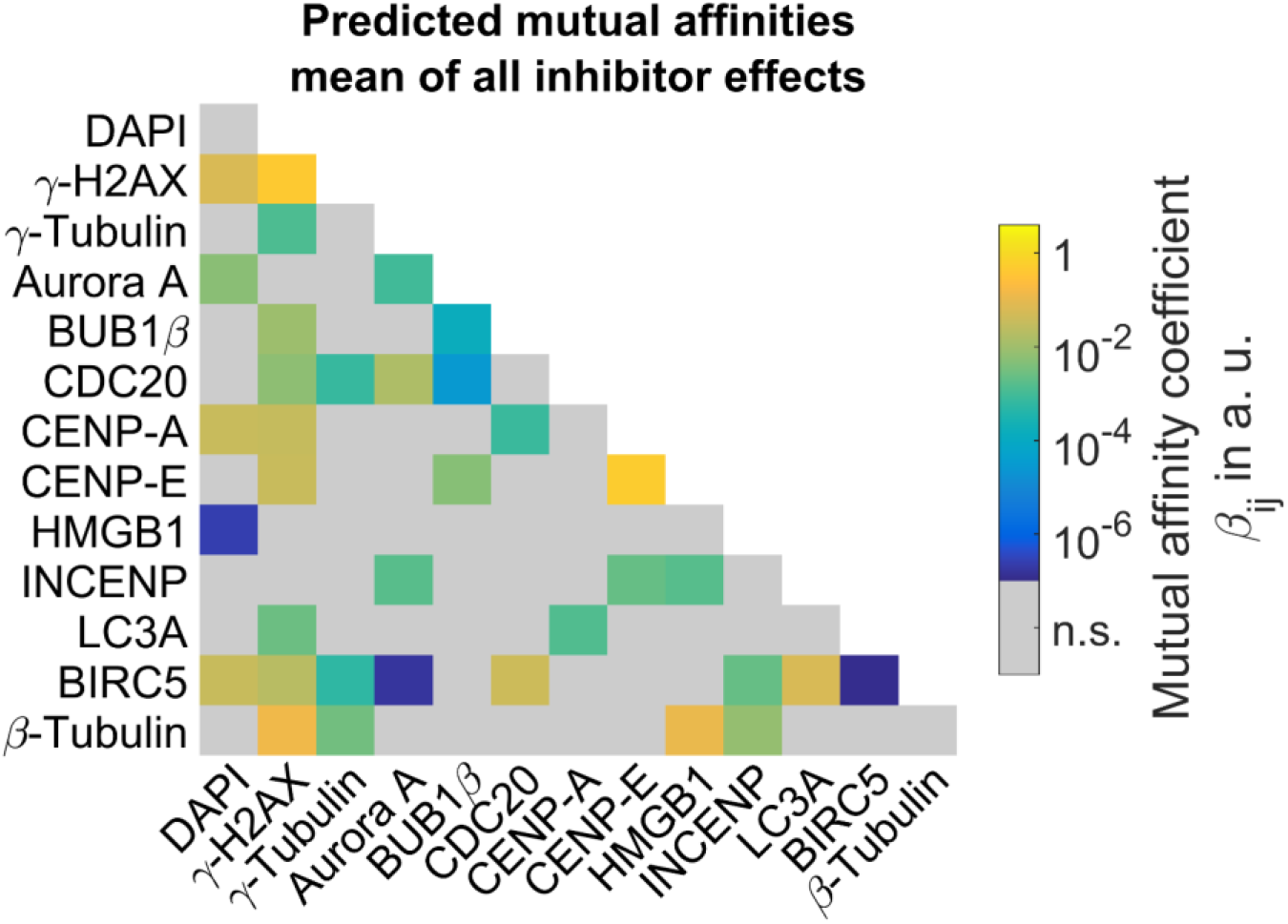
Average mutual affinities of all inhibitor effects.

**Supplementary Table 1.**
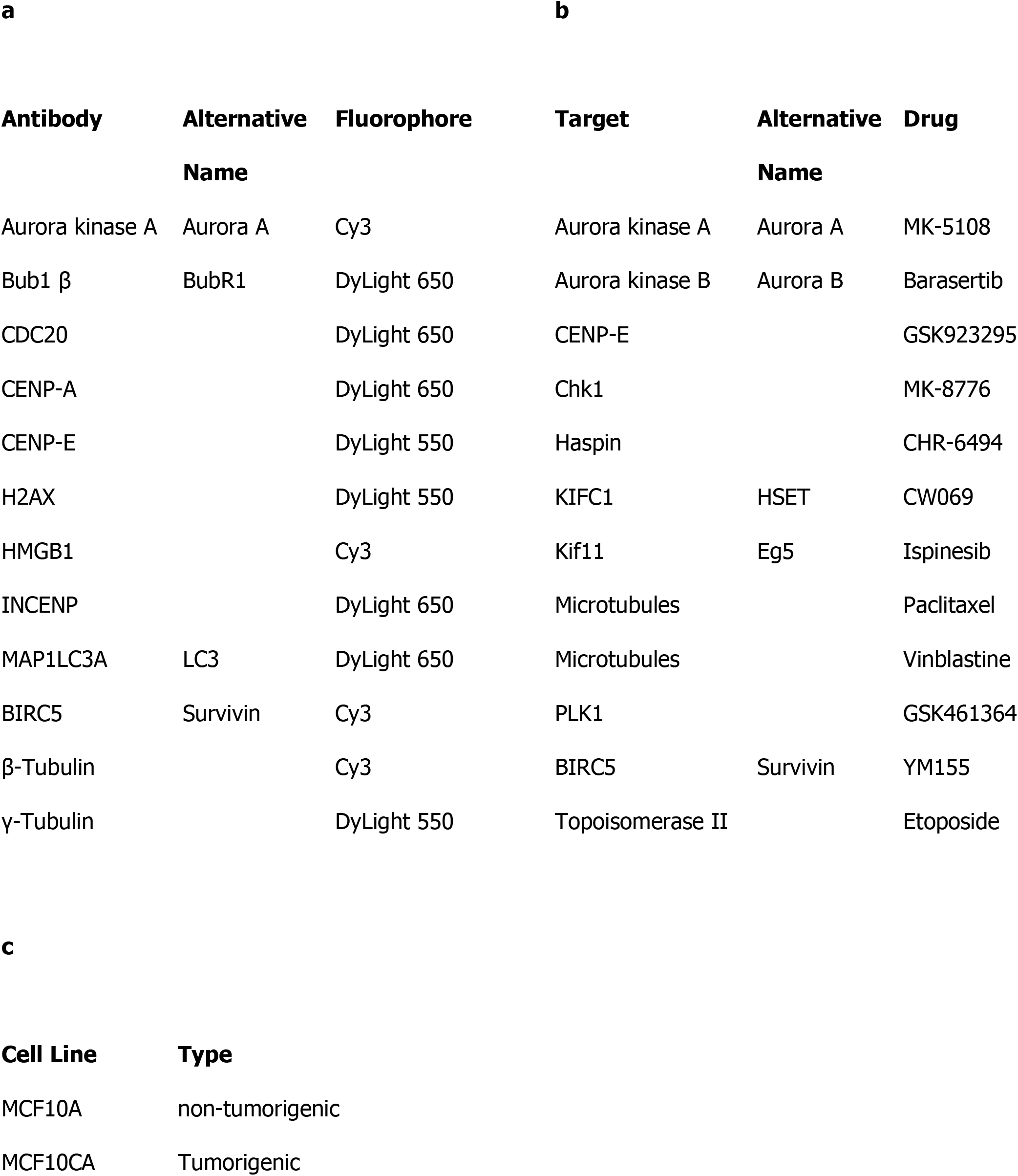
Assay proteins. (**a**) Antibodies and fluorophores. (**b**) Inhibitors and their targets. (**c**) Cell lines and their type.

| **Antibody** | **Alternative Name** | **Fluorophore** |
| --- | --- | --- |
| Aurora kinase A | Aurora A | Cy3 |
| Bub1 $\beta$ | BubR1 | DyLight 650 |
| CDC20 |  | DyLight 650 |
| CENP-A |  | DyLight 650 |
| CENP-E |  | DyLight 550 |
| H2AX |  | DyLight 550 |
| HMGB1 |  | Cy3 |
| INCENP |  | DyLight 650 |
| MAP1LC3A | LC3 | DyLight 650 |
| BIRC5 | Survivin | Cy3 |
| β-Tubulin | | Cy3 |
| γ-Tubulin | | DyLight 550 |

**b**
| **Target** | **Alternative Name** | **Drug** |
| --- | --- | --- |
| Aurora kinase A | Aurora A | MK-5108 |
| Aurora kinase B | Aurora B | Barasertib |
| CENP-E |  | GSK923295 |
| Chk1 |  | MK-8776 |
| Haspin |  | CHR-6494 |
| KIFC1 | HSET | CW069 |
| Kif11 | Eg5 | Ispinesib |
| Microtubules |  | Paclitaxel |
| Microtubules |  | Vinblastine |
| PLK1 |  | GSK461364 |
| BIRC5 | Survivin | YM155 |
| Topoisomerase II |  | Etoposide |

| **Cell Line** | **Type** |
| --- | --- |
| MCF10A | non-tumorigenic |
| MCF10CA | Tumorigenic |

### Supplementary Note 1

#### Calculation of the center of eccentricity and center of orientation measures

We compared DAPI and antibody staining intensities between cell lines, mitotic phases and for inhibitor treatments relative to controls. To this end, fluorescence intensity values were normalized by mitotic bin volumes to obtain intensity measures that were proportional to concentrations. To compare spatial distributions of protein intensities between cell lines, mitotic phases and inhibitor treatments, we defined characteristic measures of eccentricity and orientation. For each cell, 18 ROIs were defined as intersections between six eccentricity shells with indices *μ* = 1…6 and three orientations relative to the division plane (equatorial, [−30°;30°]; diagonal, [30°,60°]; polar, [60°,120°]) with indices *ν* = 1…3. The center of eccentricity *r* was defined by a sum of eccentricities *r_μ_* weighted by fluorescence intensities *I_μν_*

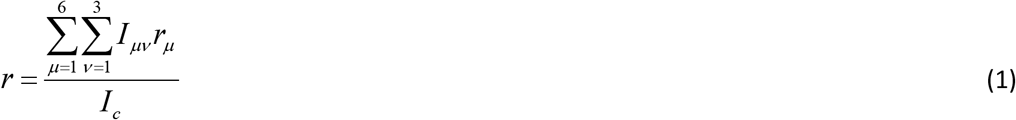

with the total sum of intensities 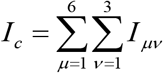. Similarly, the center of orientation *φ* was defined by

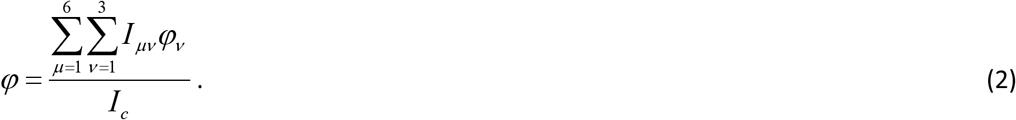

In the middle and right column of **Fig. 3a**, significant changes of *r* and *φ* dependent on cell lines, mitosis phases and inhibitor treatments were visualized. To test for significance, we performed two-sample t-tests. Given the assumption that intensity measures are in general normally distributed, this is valid for weighted sums of intensities. From applying (Bonferroni) correction for multiple testing for a total of 54 comparisons based on measurements for each protein, significance was defined by *p* < 0.05/54 ≈ 9.26 ·10^−4^.

#### Mathematical model of protein affinities to subcellular compartments and mutual affinities between proteins

To describe binding of proteins to subcellular compartments and mutual binding of proteins within subcellular compartments, we constructed a mathematical model derived from ordinary differential equations (ODEs). Here, we describe the concept and the implementation of this model.

In the simplest case, we consider binding of two proteins A_1_ and A_2_. Concentrations of free species are denoted by *A*_1_ and *A*_2_ (**Supplementary Note Figure 1**). Binding to mitotic bin *l*, results in A_1I_ and A_2I_ with concentrations *A*_1*l*_ and *A*_2*l*_. Kinetic parameters describing A_1_ binding and unbinding to this compartment are *k*_1*l*_ and *k*_−1*l*_, while A_2_ binding and unbinding is described by *k*_2*l*_ and *k*_−2*l*_. We assume that binding sites for proteins in a compartment are not limiting. The equation describing the concentration of free A_1_ thus reads

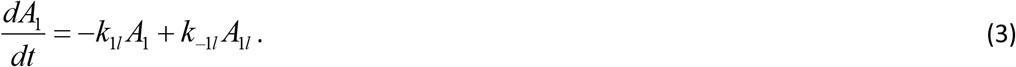

Dissociation constants *K*_1*l*_ = *k*_−1*l*_ /*k*_1*l*_ and *K*_2*l*_ = *k*_−2*l*_ /*k*_2*l*_ are thus dimensionless parameters. In steady state, the concentrations for *A*_1_ and *A*_2_ bound to mitotic bin *l* equal *A*_1*l*_ = *A*_1_ / *K*_1*l*_ and *A*_2*l*_ = *A*_2*l*_ / *K*_2*l*_.

**Supplementary Note Figure 1.**
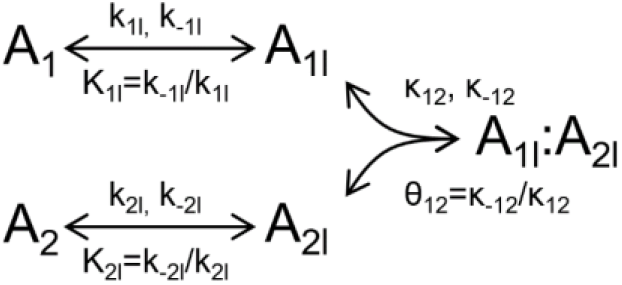
Binding of species A_1_ and A_2_ to mitotic bin *l* results in A_1I_ and A_2I_. Mutual binding in this compartment results in A_1I_:A_2I_.

In mitotic bin *l*, A_1I_ can reversibly bind to A_2I_, resulting in A_1I_:A_2I_ with concentration *A*_1*l*_ : *A*_2*l*_. We assume that binding between A_1I_ and A_2I_ depends only on the interaction between the proteins but not on the mitotic bin. Their binding and unbinding is described by parameters *㮪*_12_ and *κ*_−12_, and the dissociation constant *θ*_12_ = *κ*_−12_ / *κ*_12_. ODEs for *A*_1*l*_ and *A*_1*l*_ : *A*_2*l*_ read

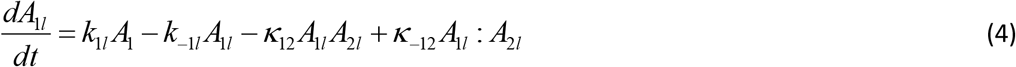

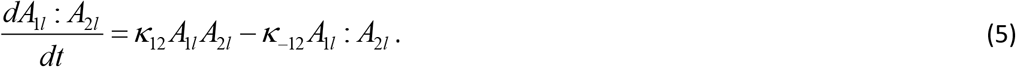

Solving at steady state for the concentration of A_1I_:A_2I_ results in

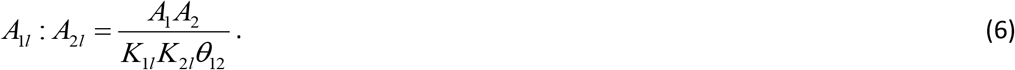

After immunofluorescence staining for A_1_, the fluorescence intensity in mitotic bin *l* is therefore given by

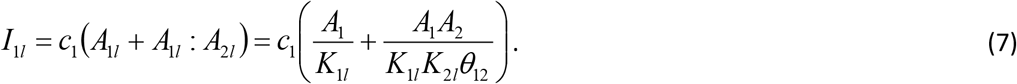

In this equation, the scaling factor *c*_1_ relates concentrations to fluorescence intensity values. The measured intensity thus depends linearly on the concentration of A_1_ and further contains the product of concentrations A_1_ and A_2_. This resembles that A_1_ and A_2_ are either recruited to mitotic bin *l* due to their affinity to this compartment or due to their mutual affinity.

To generalize this model, we describe binding of proteins A_i_ with *i* = 1…*n* to subcellular compartments *l* = 1…*m*. Affinities of proteins A_i_ to a mitotic bin are described by the n-by-m matrix *α_il_* = 1/ *K_il_*, whereas affinities between proteins are given by the n-by-n matrix *β_ij_* = 1/*θ_ij_*. Then, intensities of all species in all subcellular compartments of a cell are given by

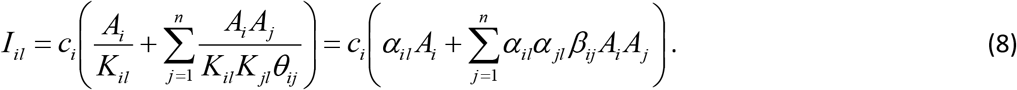

For parameter estimations, we made the simplifying assumption that the concentrations of free proteins *A_i_* were approximately proportional to the average cellular concentrations *A_i,t_* = *s_i_A_i_*, with the proportionality factor *s_i_* ≥ 1. This assumption holds true if the proteins are recruited to subcellular compartments due to their direct affinities to mitotic compartments rather than their affinities to the other observed proteins, which will be justified below. Average cellular intensities were calculated by weighting intensities in subcellular compartments *I_il_* with mitotic bin volumes *V_l_*, given by

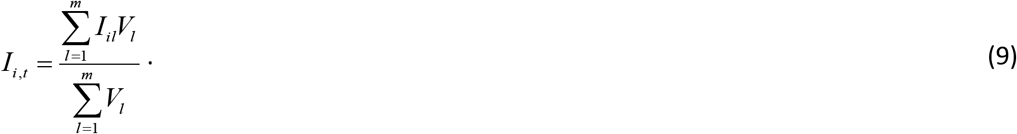

For model fitting, we calculated fold changes relative to medians of average concentrations for the population of cells, *Î_i,t_*, for intensities in subcellular compartments, *Ĩ_il_* = *I_il_* / *Î_i,t_*, and for average cellular concentrations, *Ĩ_i,t_* = *I_i,t_* / *Î_i,t_*. Then, experimental measurements could be related to model variables *A_i_* by

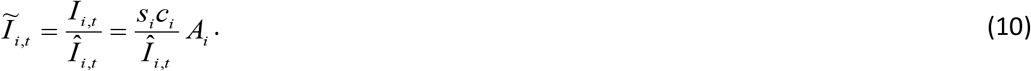

Thereby, Eq. 8 was rescaled to

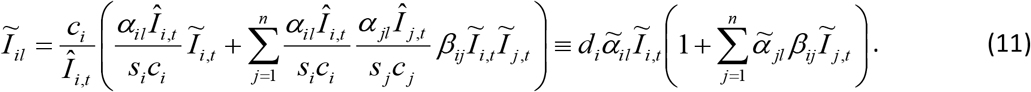

Therein, the rescaled parameter *d_i_* = *c_i_* / *Î_i,t_* was equal to the inverse of the median average cellular concentration, and 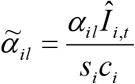 was equal to the median concentration of species A_i_ that was bound in mitotic bin *l*.

To account for different affinities of proteins to subcellular compartments during metaphase and segregation, 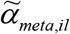 and 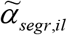, as well as scaling factors *d_meta,i_* and *d_segr,i_*, intensity measurements during these cell cycle phases were separately described by

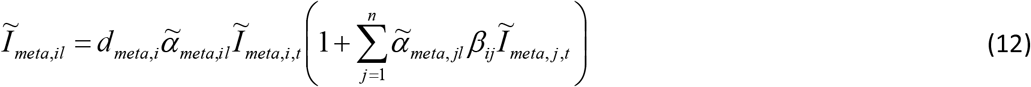

and

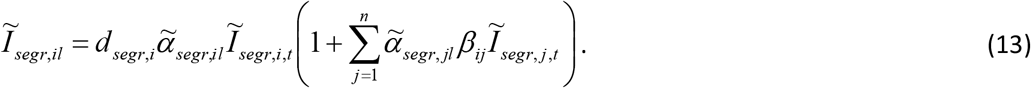

Thereby, we assume that rescaled affinities to subcellular compartments, but not mutual affinities between proteins *β_ij_*, were dependent on the cell cycle phase. We simultaneously fitted Eqs. (12) and (13) to experimental data for estimating *d_meta,i_*, *d_segr,i_*, 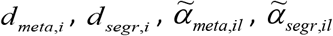 and *β_ij_*.

To analyze relevant interactions between proteins, we first used experimental data from controls (cells not treated with inhibitors). Data from MCF10A and MCF10CA cells were fitted together, assuming that differences between cell lines were only dependent on different average cellular concentrations of all proteins but not on affinities to subcellular compartments or affinities between proteins. A total of 513 to 529 parameters were estimated by model fitting to 50.635 data points (47.970 intensity measurements for subcellular compartments and 2665 average intensity values) from control experiments in 205 cells. To equally weight residuals for data points of different magnitudes we assumed the error model

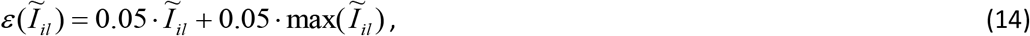

assuming that for each measurement the experimental error is given by 5% of the measurement value plus 5% of the maximal value of all included cells. For model fitting, we used the MATLAB solver lsqnonlin, to minimize residuals between measurements

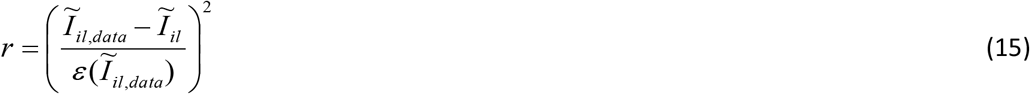

for all cells. After initial fittings, parameter intervals were defined for 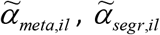, *d_meta,i_* and *d_segr,i_* between 10^−3^ and 10^2^, and for *β_ij_* between 10^−7^ and 10^2^. To accelerate convergence of model fits, parameters were fitted on a log-scale.

First, we started with fitting a model that only accounts for known literature interactions that were extracted from the *Ingenuity* pathway knowledge (IPA) database. To this end, *β_ij_* was reduced to entries according to this set of literature interactions regarded as ground truth. Then, by sequential feature selection, additional affinities between proteins were further included if they could significantly improve the squared sum of residuals of the model fit. For each selection step of testing whether an additional interaction should be included, we performed 50 multi-start local optimizations by sampling initial conditions from allowed parameter intervals. We assured that after optimizations, differences between the best fits were below the residuals for single data points. Additional entries in *β_ij_* were selected based on likelihood-ratio testing, assuming that the likelihood-ratio for a model including an additional variable compared to a model without the additional parameter follows a one-dimensional *χ*^2^ distribution. An additional affinity between proteins was included in *β_ij_* if the increase in log-likelihood exceeded the 95% confidence interval of the cumulative one-dimensional *χ*^2^ distribution. Following this forward selection procedure, 16 additional affinities between proteins were included. The reduction of the residual sum of squares is shown in **Supplementary Note Figure 2**.

**Supplementary Note Figure 2.**
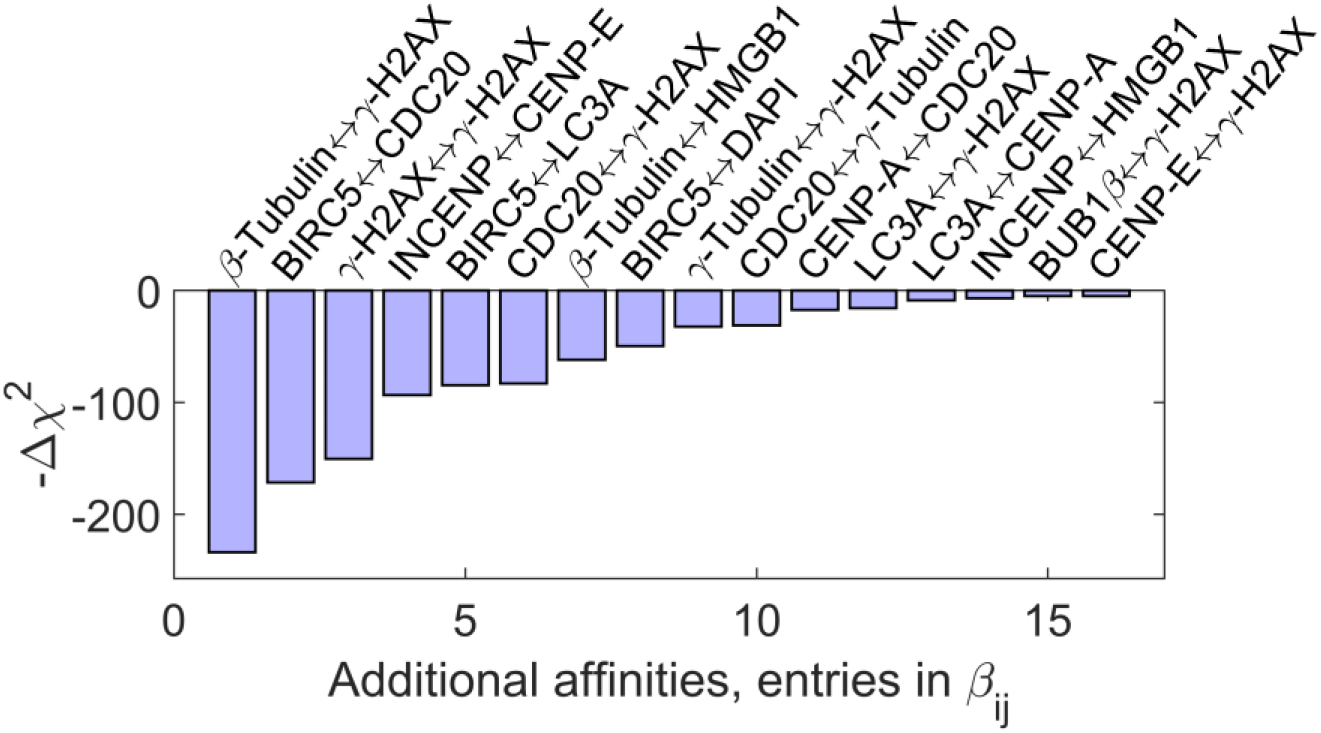
Affinities between proteins included in addition to literature interactions. By sequential forward selection, entries in the matrix *β_ij_* that contains mutual affinities between proteins were included in the model if the model fit was significantly improved. All included affinities were ordered according to the improvement in the *χ*^2^ measure of model deviation from the experimental data.

These additional entries in *β_ij_* represent hypotheses about mutual binding between proteins. Notably, this predicted mutual binding may be distinct from possible functional interactions between proteins. Mutual affinity does not necessarily imply a functional interaction, whereas a functional interaction may not require high binding affinity. Nevertheless, predictions of mutual affinities between the observed proteins involved in mitosis can be used to guide further experiments for investigating functional relations and protein complexes that are linked to cellular processes.

After identifying an optimal set of additionally included affinities from model fitting to the control dataset, the model was fitted to data from inhibitor treatments. For every inhibitor treatment, the parameters *d_meta,i_*, *d_segr,i_*, 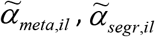 and the extended *β_ij_* were estimated by model fitting.

**Supplementary Note Figure 3.**
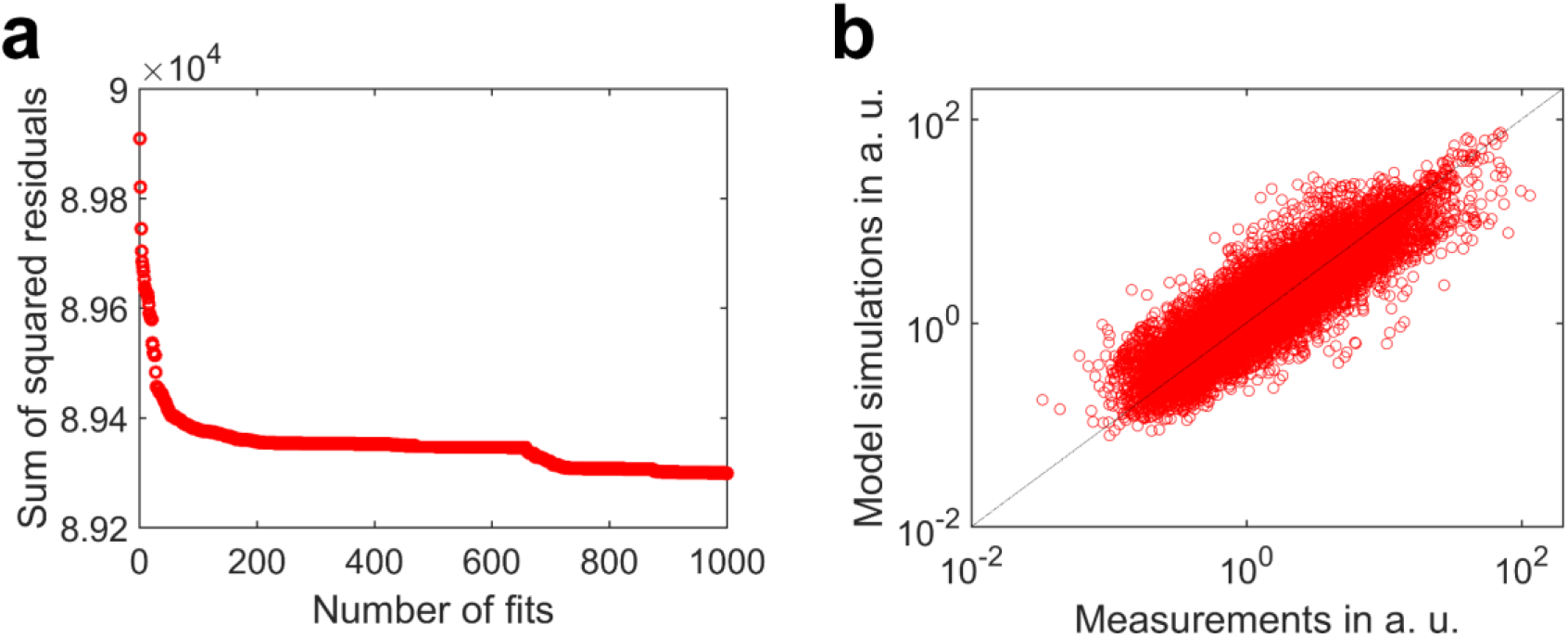
Results of multi-start local optimizations. (**a**) Ordered sums of squared residuals for 1000 model fits to the control dataset from untreated cells. Differences between best fits were below the range of squared residuals for single data points indicating convergence of model fits towards a global optimum. (**b**) Best-fit model simulations and measurements indicate that the model is consistent with the experimental dataset.

Finally, to estimate effects from inhibitor treatments on compartment affinities and on mutual affinities between proteins, we again fitted the model to the control dataset from untreated cells and to datasets from inhibitor treatments. We performed in each case 1000 multi-start local optimizations by sampling initial conditions from allowed parameter intervals. **Supplementary Note Figure** shows the ordered sum of squared residual values for 1000 multi-start local optimization runs for fitting the control dataset. In **Supplementary Note Figure 3b** model simulations for the best model fit to the control dataset were plotted against experimental data. It is evident that the model fit is highly consistent with experimental measurements. Known and additionally predicted entries of *β_ij_*, and estimated affinity values for the control dataset were shown in **Fig. 3c** and **3d**. Estimated affinity values for an exemplary inhibitor (PLK1) were visualized in **Fig. 3e**. Furthermore, the average affinities for all inhibitors were shown in **Supplementary Fig. 2**. **Supplementary Fig 2c** shows estimates 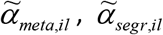 for estimated affinities to subcellular compartments, whereas **Supplementary Fig 2d** shows estimates *α_meta,il_* / *s_i_* and *α_segr,il_* / *s_i_* that were obtained by multiplying with scaling factors *d_meta,i_* and *d_segr,i_*.

